# Reconstruction of the cell pseudo-space from single-cell RNA sequencing data with scSpace

**DOI:** 10.1101/2022.05.07.491043

**Authors:** Jie Liao, Jingyang Qian, Ziqi Liu, Ying Chi, Yanrong Zheng, Xin Shao, Junyun Cheng, Yongjin Cui, Wenbo Guo, Penghui Yang, Yining Hu, Hudong Bao, Qian Chen, Mingxiao Li, Bing Zhang, Xiaohui Fan

**Author notes:** Corresponding Authors: Prof. Xiaohui Fan, and Dr. Bing Zhang. These authors contribute equally to this work (Jie Liao, Jingyang Qian, Ziqi Liu, Ying Chi).

## Abstract

Tissues are highly complicated with spatial heterogeneity in gene expression. However, the cutting-edge single-cell RNA-seq technology eliminates the spatial information of individual cells, which contributes to the characterization of cell identities. Herein, we propose **<u>s</u>**ingle-**<u>c</u>**ell **<u>s</u>**patial **<u>p</u>**osition **<u>a</u>**ssociated **<u>c</u>**o-**<u>e</u>**mbeddings (scSpace), an integrative algorithm to distinguish spatially variable cell subclusters by reconstructing cells onto a pseudo-space with spatial transcriptome references (Visium, STARmap, Slide-seq, etc.). We demonstrated that scSpace can define biologically meaningful cell subpopulations neglected by single-cell RNA-seq or spatially resolved transcriptomics. The use of scSpace to uncover the spatial association within single-cell data, reproduced, the hierarchical distribution of cells in the brain cortex and liver lobules, and the regional variation of cells in heart ventricles and the intestinal villus. scSpace identified cell subclusters in intratelencephalic neurons, which were confirmed by their biomarkers. The application of scSpace in melanoma and Covid-19 exhibited a broad prospect in the discovery of spatial therapeutic markers.

## Introduction

Uncovering the organization of cells in a tissue and how this organization affects function is a fundamental pursuit of life science research^1^. Tissue complexity is defined by the diversity of cell types and spatial heterogeneity of gene expression^2-4^. The emergence of single-cell RNA-seq^5-8^ and spatial transcriptomics technologies^9-13^ can resolve the structural and functional partitioning of tissues from the cellular and spatial dimensions, respectively. However, there seems to be a natural gap between these two technologies. A classic protocol of scRNA-seq is to digest tissues into single-cell suspension, which leads to the loss of spatial information of cells. Meanwhile, the state-of-the-art spatially resolved RNA-seq technologies could, though, capture the unbiased transcriptome of each spot, fail to achieve cellular resolution^14-16^. Although *In silico* methods have made great progress in spot deconvolution^17-19^, spatial mapping^20, 21^, and gene imputation^22-24^ via integrative analysis of single-cell and spatially resolved transcriptomics. Few approaches are reported to use spatial transcriptomics to conduct more refined cell classification of single-cell transcriptomes despite that in many circumstances, spatial characteristics of tissues have proved to be a key determinant in cell identification and functioning^25, 26^. Consequently, in addition to transcriptomes, spatial relationships among cells should also be taken into consideration while delineating the cellular microenvironment^27^ and resolving the cell-cell communication^28-30^.

To this end, we introduce scSpace, a comprehensive algorithm that integrates spatially resolved transcriptomics data to reconstruct spatial associations of single cells within scRNA-seq data (Extended Data Fig. 1a). Using a transfer learning model, termed Transfer Component Analysis (TCA), scSpace could extract the characteristic matrixes from both spatial transcriptomics and scRNA-seq data, and project single cells into a pseudo-space via a multi-layer neural network (MNN) model, so that gene expression graph and spatial graph of cells can be embedded jointly for the further spatial reconstruction and space-informed cell clustering with higher accuracy and precision. Moreover, both simulated datasets and existing spatial transcriptomics data were utilized to benchmark the performance of scSpace. Finally, we validated scSpace with regional single-cell data and then applied it to discover significant cell subpopulations with spatial heterogeneity as well as transcriptional specificities in diseases, such as melanoma and Covid-19, at single-cell resolution.

## Result

### Design concept and performance evaluation of scSpace on simulated data

Although the mainstream spatial transcriptomics^10, 11^ was obtained by sequencing spots containing a mixture of cells, whereas scRNA-seq labels individual cells before pooling and sequencing, both methods share comparable gene expression and transcriptional states^4^. Therefore, in this study, a transfer learning model was used to integrate single-cell data *X*_*S*_ and spatial transcriptomics data *X*_*T*_, preserving both the transcriptional information and the spatial relationships of cells. As illustrated in Fig. 1a and Extended Data Fig. 1a, scSpace first utilized TCA to eliminate the batch effect of these two types of data and extract a latent feature representation *ϕ* across both domains with true biological characteristics. After the transformation was determined, single cells were mapped to an assumed coordinate system (pseudo-space) characterized by the MNN model using existing spatial references such as Visium, STARmap, ST, Slide-seq, etc. The resulting cells in the pseudo-space contain an expression graph *G*_*g*_(*V, E*_1_) and a space graph *G*_*S*_(*V, E*_2_), which were constructed using the *k*-nearest neighbor algorithm. Then, cells were clustered to identify spatially heterogeneous cell subpopulations based on the space-informed expression graph which was retrieved by transforming *G*_*S*_(*V, E*_2_) to a spatial weight *w* of each edge in *G*_*g*_(*V, E*_1_).

**Fig. 1.**
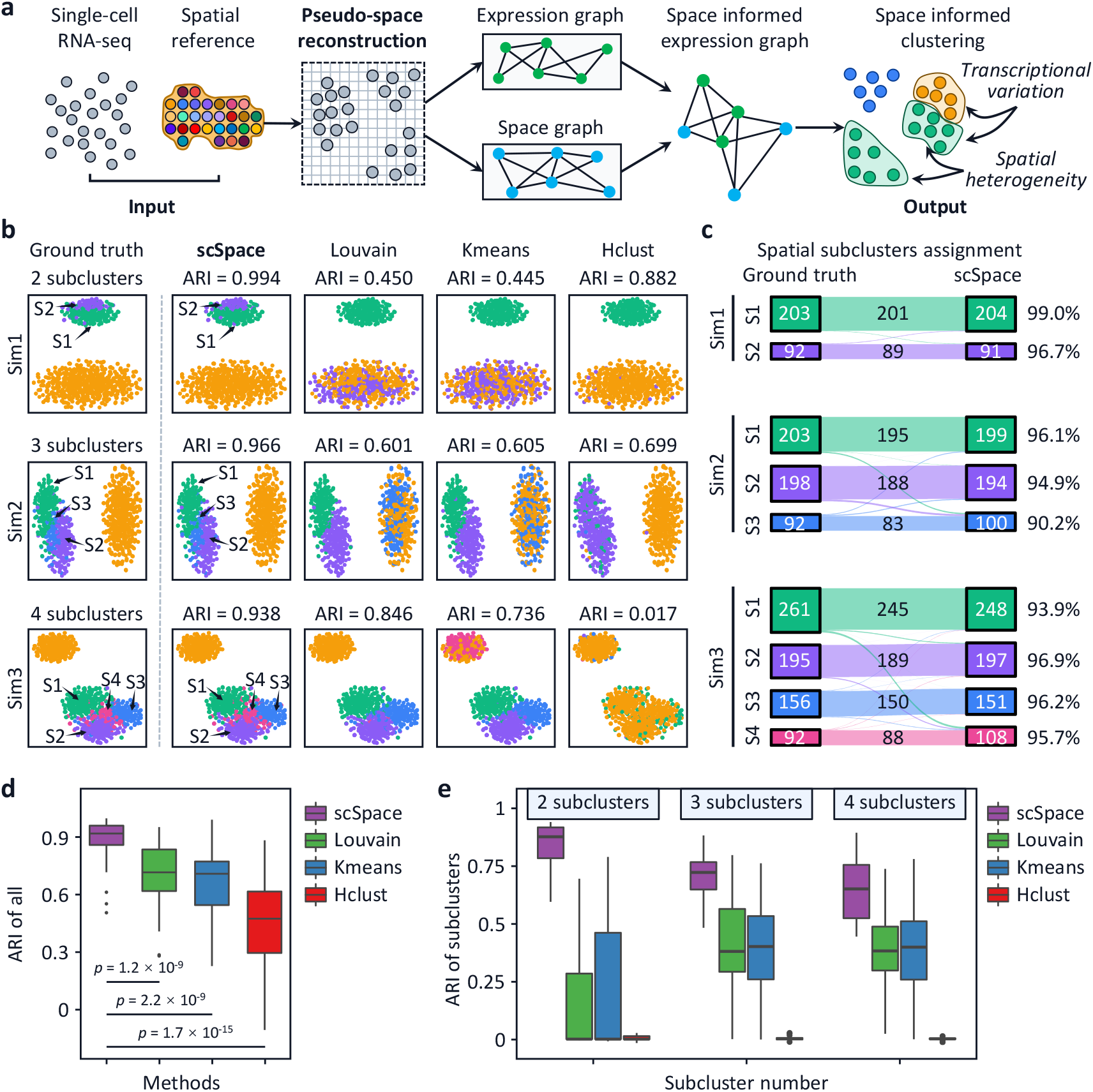
Schematic workflow of scSpace and performance evaluations on simulated data. **a**, Overview of the design concept of scSpace. First, single-cell and spatial reference data (input) were co-embedded to reconstruct the pseudo-space for individual cells, which contained an expression graph and a space graph. Second, the space graph was transformed to edge weights in the expression graph to construct a space-informed expression graph. A space-informed cell clustering (Output) was performed to uncover both transcriptionally variated and spatially heterogeneous subpopulations. **b**, Comparison of different clustering methods (scSpace, Louvain, Kmeans, and Hclust) using simulated datasets containing different numbers of clusters and subclusters (Three examples of simulated ground truth data with two, three, and four subclusters, respectively). The ARI score was introduced to evaluate the accuracy of the clustering performance for each method. Sn devotes Subcluster n to the simulated data. **c**, Jaccard index demonstrated the number of cells that were successfully assigned by scSpace to their corresponding subclusters in the ground truth. **d**, Performance of scSpace compared with Louvain, Kmeans, and Hclust on 50 simulated data using ARI scores of all clusters and subclusters. (the average ARI score of 0.914 for scSpace was significantly higher than Louvain, Kmeans, and Hclust with *p*=1.2×10^−9^, *p*=2.2×10^−9^, *p*=1.7×10^−15^, respectively). **e**, ARI scores of only spatially variated subclusters (from 2 to 4) across all 50 simulated datasets for scSpace and other methods.

Next, we benchmarked scSpace with classical cell clustering methods, Louvain, K-means (Kmeans), and Hierarchical clustering (Hclust) using 50 simulated data which were constituted with different spatial distribution patterns including several clusters and 2 to 4 refined subclusters (Extended Data Fig. 1b). Compared with other methods, scSpace could distinguish spatially variated subclusters in acceptable agreement with the ground truth with higher adjusted rand index (ARI) scores (Fig. 1b and Extended Data Table S1). Also, scSpace gained higher accuracy of cell assignment for each spatial subcluster in the ground truth (Fig. 1c). As shown, scSpace performed superior to Louvain, Kmeans, and Hclust with significantly higher ARI scores in distinguishing all clusters (Fig. 1d), and only spatially heterogeneous subclusters (Fig. 1e). As the subcluster number of cells increased in simulations, the Pearson correlation between the pseudo-space and the ground truth decreased slightly (Extended Data Table S2). Overall, scSpace reconstructed the pseudo-space of single cells with high correlations in the distances to original positions (Extended Data Fig. 1c, and d).

### Reconstruction of the hierarchical structure of human and mouse cortex using biological data by scSpace

Spatially resolved transcriptomics data are better examples than simulated data for evaluating the reconstruction performance of scSpace because cell coordinates in spatial data are biologically meaningful. Thus, we first collected spatial transcriptomics data of human dorsolateral prefrontal cortex (DLPFC)^31^ and mouse primary visual cortex (V1)^9^, which were profiled by Visium and STARmap, respectively. Second, for each set of spatial transcriptomics data, we selected one slice as the spatial reference, and another slice with cell coordinates removed as test data.

As illustrated in Fig. 2a, scSpace successfully reconstructed the hierarchical structure of different human DLPFC layers in the pseudo-space, with the relative position between the layers preserved. The Normalized pairwise distances between spots in the pseudo-space were calculated and found to increase monotonically with the pairwise distances in the original tissue, which is consistent with our structural correspondence assumption (Fig. 2b). Further analysis demonstrated that the pairwise distances between spots in the embedded pseudo-space and the ground truth were highly correlated (Fig. 2c). Moreover, we examined the spatial distribution of marker genes (HPCAL1, HOPX, MEFH, PCP4, KRT17, and MOBP) in different layers (Layer 2, Layer 3, Layer 4, Layer 5, Layer 6, and WM, respectively), and found that scSpace predicted gene expression patterns correlated well with the original patterns (Fig. 2d). More examples used to validate the performance of scSpace were shown in the Extended Data Fig. 2a. Similar results were reproduced when we tested scSpace using another spatially resolved single-cell transcriptomics data with 1020 targeted genes, which were derived from mouse V1 neocortex by STARmap (Fig. 2e and Fig. 2f).

**Fig. 2.**
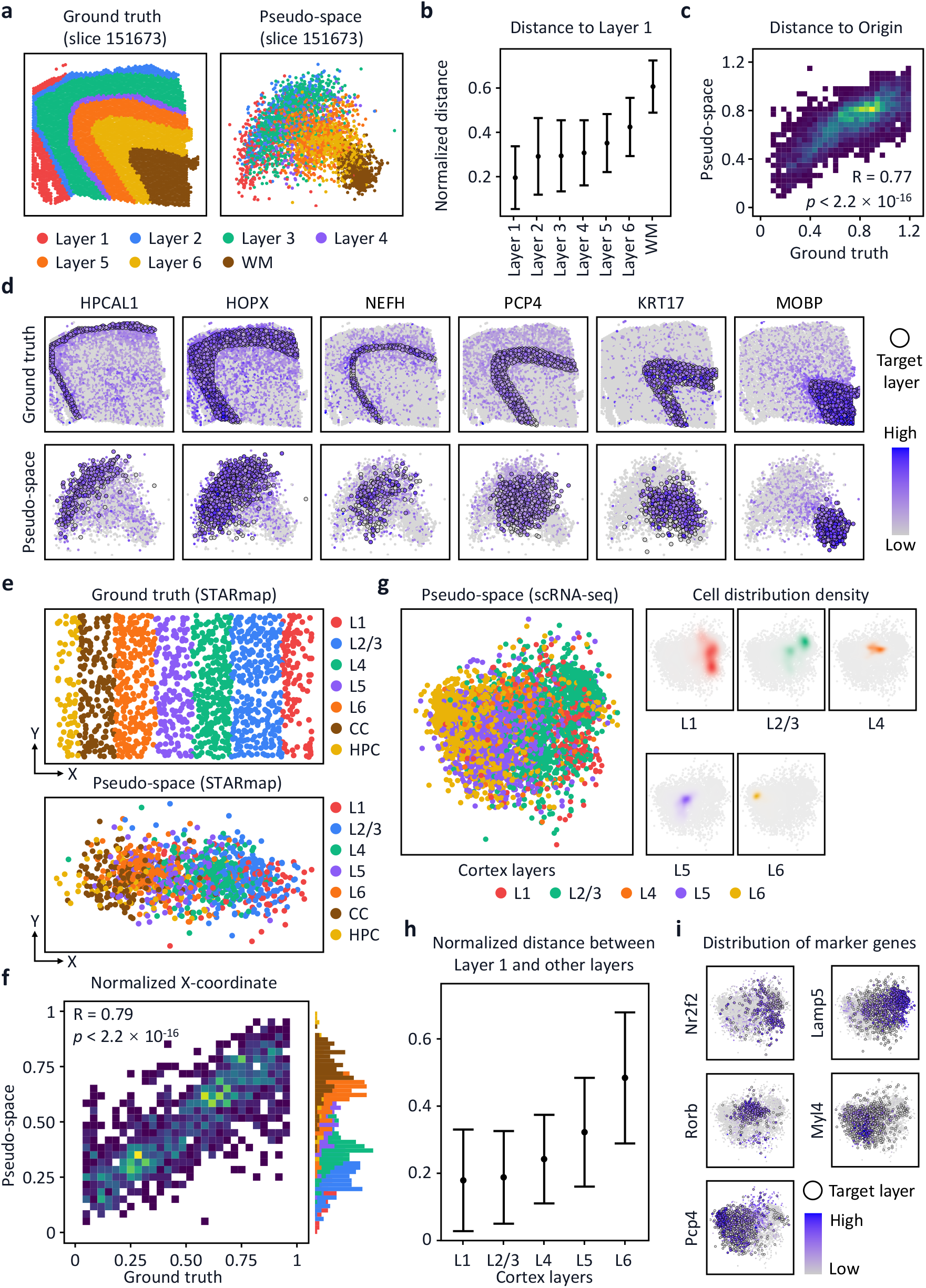
Reconstructing cortex hierarchical structure using spatial transcriptomics data and validation on single-cell RNA-seq data. **a**, Spatial reconstruction of human DLPFC layers using one slice as the spatial reference and another as the test set by removing the cell coordinate information. **b**, Normalized pairwise distances between spots from different layers and Layer 1. **c**, Correlation of scSpace predicted spot coordinates and their original locations. **d**, Gene expression of layer landmarks in the spatial transcriptomics and the pseudo-space. **e**, Spatial reconstruction of mouse V1 neocortex layers using STARmap data. **f**, Correlation of scSpace predicted cell coordinates and their original locations. **g**, Pseudo-space (Left) and distribution pattern of cells in each layer (Right) of the scRNA-seq data predicted by scSpace. **h**, Normalized pairwise distances between cells from different layers and Layer 1. **i**, Marker gene expression patterns of cells from different layers in the predicted pseudo-space.

Next, we applied scSpace to reconstruct the pseudo-space of a real scRNA-seq dataset derived from Allen Brain Atlas (see Methods), the results confirmed the hierarchical structure of the mouse V1 neocortex. The spatial distribution pattern of each layer was different and the relative positions between these layers corresponded agreeably to the spatial reference (Fig. 2g). The pairwise distances between single cells from different layers and Layer 1 in the pseudo-space and the physical space increased synchronously (Fig. 2h). The spatial gene expression pattern of the biomarkers (Nr2f2, Rorb, Pcp4, Lamp5, and Myl4) correlated well with the spatial distribution of individual cells from corresponding layers in the pseudo-space (Fig. 2i).

### Regional reconstruction of scRNA-seq data in different circumstances using scSpace

To investigate the ability of scSpace in restoring the relative spatial associations among cells, we focused on real scRNA-seq data that were obtained from seven distinct zones of an intestine^32^ (Fig. 3a), nine layers of a liver lobule^33^ (Extended Data Fig. 3a), and two segregated regions of an embryonic heart^34^ (Fig. 3d), based on the robust marker gene expression. After the cells were allocated to a pseudo-space, we found that the distribution of cells varied from region to region (Fig. 3b and Extended Data Fig. 3b, d), exhibiting a diverse functioning zonation in the tissue microenvironment. Along with the intestinal villus axis from V1 to V6, the pairwise distances between cells to the villus crypt showed an increasing trend (Fig. 3c), which was consistent with the expectation. Similar results were reproduced on the regional reconstruction of the liver lobule (Extended Data Fig. 3c) and embryonic heart. We further validated the expression patterns of marker genes in different regions.

**Fig. 3.**
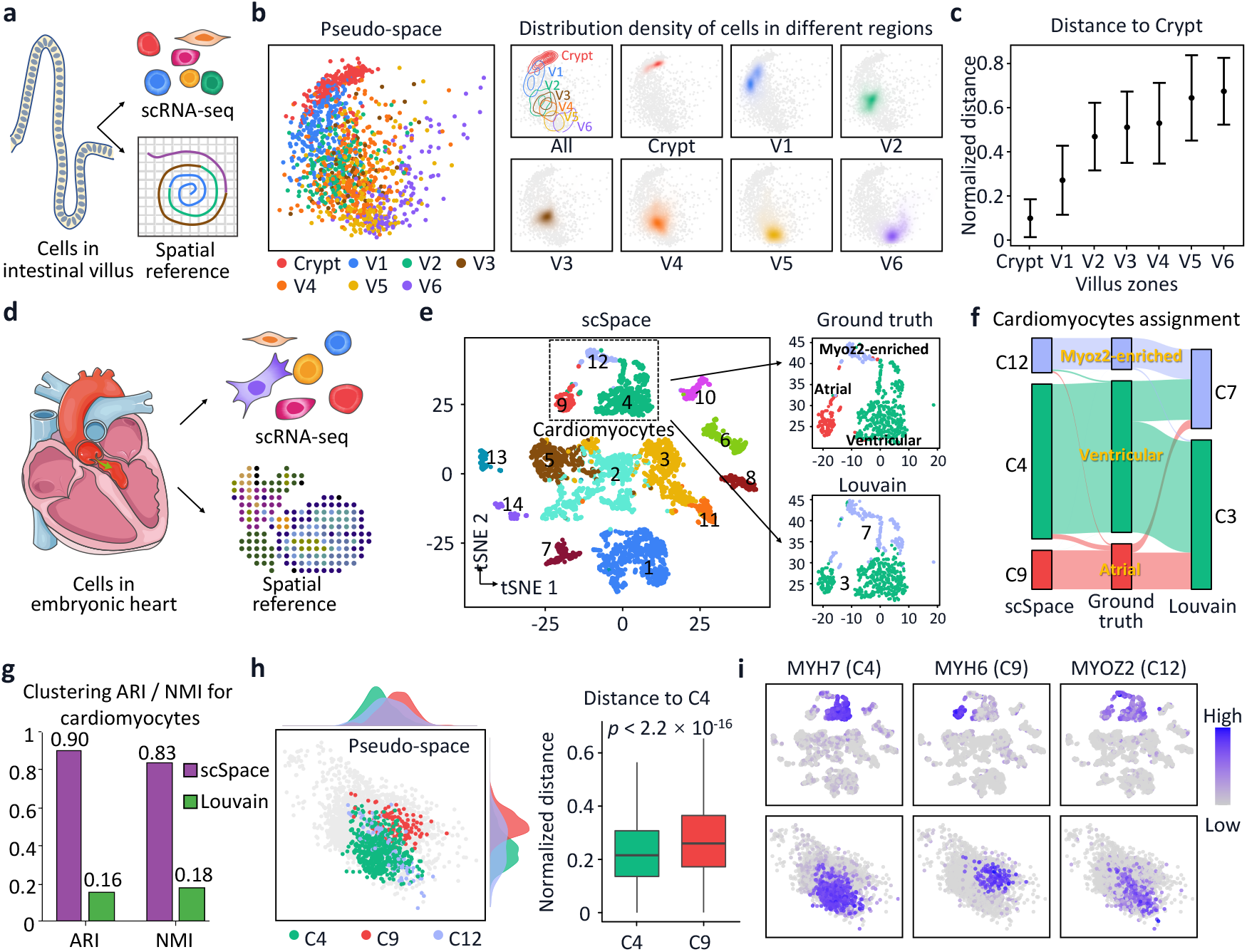
Validation of scSpace on regional scRNA-seq data. **a**, Schematic of the spatial reconstruction of intestinal epithelium using biological scRNA-seq data from different regions. Both the single-cell data and spatial reference were accessed from experimental data. **b**, Left, the pseudo-space of intestinal cells generated by scSpace. Right, the distribution densities of cells in different regions (All, Crypt, V1, V2, V3, V4, V5, and V6). **c**, Normalized pairwise distances between cells in different zones and cells in the villus crypt from the near to the distant. **d**, Spatial reconstruction of atrial, ventricular, and Myoz2-enriched cardiomyocytes in the embryonic heart with real scRNA-seq data and spatially resolved transcriptomics. **e**, Space-informed clustering of cells in the embryonic heart using scSpace. Left, refined clustering of the original data obtained 14 subpopulations. Right, three subpopulations of cardiomyocytes C4, C9, and C12 predicted by scSpace correspond to ventricular, atrial, and Myoz2-enriched cardiomyocytes annotated in the ground truth, respectively. The Louvain algorithm classified cardiomyocytes into two subclusters. **f**, Comparison of the number of cells scSpace and Louvain successfully assigned to their corresponding subclusters in the ground truth using the Jaccard index. **g**, Evaluation of the clustering performance of scSpace and Louvain using ARI and NMI scores. scSpace, ARI=0.90 and NMI=0.83, Louvain, ARI=0.16 and NMI=0.83. **h**, Spatial distribution of three cardiomyocyte subpopulations C4, C9, and C12 in the pseudo-space. Left, the pseudo-space generated by scSpace and the distribution of C4 (green), C9 (red), and C12 (light indigo) along the X- and Y-axis of the pseudo-space. Right, pairwise distances between cells in ventricular (C4) and cells in atrial (C9) subpopulations. **i**, Expression patterns of marker genes for C4, C9, and C12 subpopulations in the pseudo-space. MYH7 for C4, MYH6 for C9, and MYOZ2 for C12, respectively.

Specifically, clustering analysis of scRNA-seq data of the embryonic heart was carried out using scSpace and Louvain, and then compared with the original annotations provided experimentally. As illustrated in Fig. 3e and Fig. 3f, atrial (C9), ventricular (C4), and Myoz2-enriched (C12) cardiomyocytes can be precisely distinguished by scSpace, which was confirmed by the experimental data. However, the Louvain algorithm missed the spatial variation within the single-cell data. The clustering accuracy (ARI and normalized mutual information (NMI) scores) of scSpace were 0,90 and 0.83, respectively, which were significantly higher than that of Louvain (0.16 and 0.18), as shown in Fig. 3g. The distribution patterns of the three subpopulations of cardiomyocytes in the pseudo-space were segregated, and the normalized pairwise distances between atrial (C9) and ventricular (C4) cardiomyocytes were significantly distinguished from each other (Fig. 3h). The spatial expression patterns of relevant marker genes for C9, C4, and C12 subpopulations were also significantly correlated with the distribution of cells (Fig. 3i and Extended Data Fig. 3e). The application in real single-cell sequencing data indicated that scSpace has great potential to distinguish spatially variated cell subclusters.

### Discovery of spatially variated subpopulations in human cortex from scRNA-seq data

As mentioned above, scSpace can preserve the spatial associations of human DLPFC cells by reconstructing the pseudo-space using de-coordinated spatial transcriptomics data. Here we further demonstrate its ability on deciphering spatially variated subpopulations in the human cortex from experimental scRNA-seq data accessed from Allen Brain Atlas (see Methods). As shown in Fig. 4a, scSpace first embedded single cells in a pseudo-space to reconstruct the spatial associations of cells. Consistent with previous results, after spatial reconstruction by scSpace, the distribution density of cells in different cortex layers was significantly different in the pseudo-space (Fig. 4b) and the normalized pairwise distances between cells and Layer 1 (L1) increased layer by layer from L1 to WM (Fig. 4c). The results showed that the pseudo-space reconstructed by scSpace has biological significance and rationalized the subsequent space-informed clustering based on it.

**Fig. 4.**
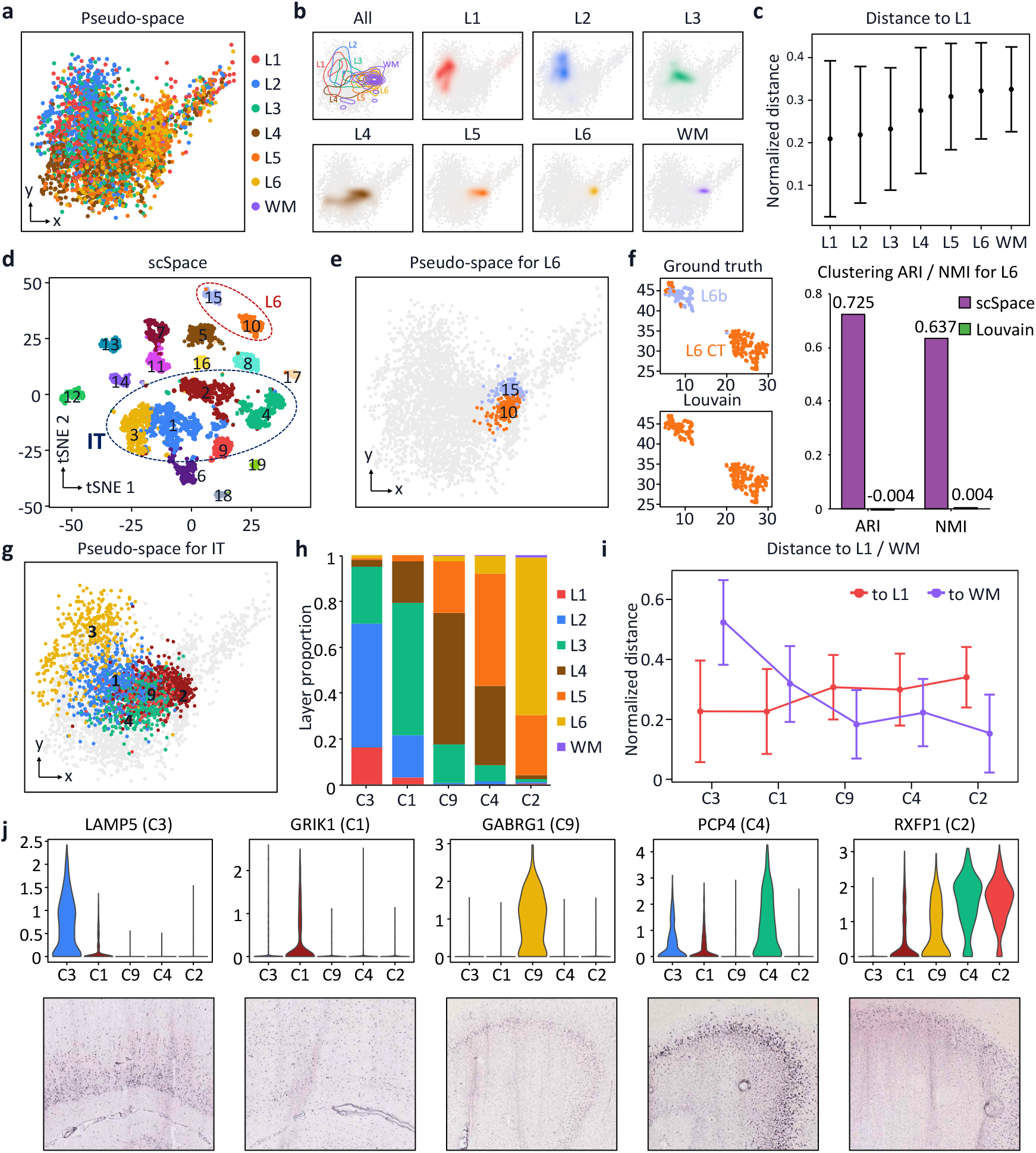
Discovery of fine subpopulations in human cortex by scSpace. **a**, Spatial reconstruction of human cortex cells. **b**, Distribution densities of cells in different cortex layers. **c**, Normalized pairwise distances between different cells from different layers and Layer 1. **d**, Space-informed clustering analysis of scRNA-seq to identify refined cell subpopulations using scSpace. **e**, Spatial distribution of the two subclusters with spatial heterogeneity C10 and C15, predicted by scSpace. **f**, Left, annotation of each cell in C10 and C15 subclusters in the ground truth (Top) and transcriptome-based clustering results analyzed by Louvain (Bottom). Right, clustering accuracy (ARI and NMI scores) of scSpace (purple) and Louvain (green) in distinguishing cells from C10 and C15 subpopulations. **g**, Cellular pseudo-space reconstruction of refined IT subclusters by scSpace. **h**, The distribution proportion of each cell subpopulation on each cortex layer. **i**, Normalized pairwise distances between IT neurons from L1 to WM, and vice versa. **j**, Expression value of marker genes of different subtypes (Top) and validation of their expression patterns on Allen Brain Atlas (Bottom).

Next, by combining transcriptional and spatial information of single cells, scSpace classified human DLPFC cells into 19 refined subpopulations (Fig. 4d). Among these subclusters, there were two subclusters in Layer 6 (L6), cluster 10# (C10) and cluster 15# (C15), exhibiting diverse spatial patterns in the pseudo-space (Fig. 4e). However, the spatial heterogeneity was difficult to distinguish by traditional algorithms such as Louvain, which uses only transcriptional information with a clustering ARI score of - 0.004 and an NMI score of 0.004, compared with that of 0.725 and 0.637 for scSpace, respectively (Fig. 4f). Once again, the results confirmed that the spatial information of each cell is crucial for the characterization of its cellular identity.

Notably, as illustrated in Fig. 4d, intratelencephalic (IT) neurons in the original dataset can be further classified into five subclusters (C3, C1, C9, C4, and C2) based on their spatial characteristics (Fig. 4g). scSpace analysis showed that IT neurons were distributed in all layers but accounted for different proportions (Fig. 4h). Moreover, the density centers of cell spatial distribution in C3, C1, C9, C4, and C2 moved from cortex L1 to WM gradually (Fig. 4i). Five genes were selected from all differentially expressed marker genes for the subpopulations (Extended Data Fig. 4a), LAMP5 (C3), GRIK1 (C1), GABRG1 (C9), PCP4 (C4), and RXFP1 (C2), were validated by the histological staining images derived from the Allen Brain Atlas. As shown, the spatial expression patterns of these marker genes were consistent with the distribution of the corresponding subclusters and exhibited a hierarchical structure (Fig. 4j).

### Spatial reconstruction of T cell pseudo-space revealed T cell exhaustion in melanoma

After comprehensive evaluations of the performance of scSpace with simulated and biological data, we seek for deeper exploration of its application prospect. We first applied scSpace to melanoma, a cancer disease with high spatial heterogeneity, to reconstruct the pseudo-space of immune cells in the tumor microenvironment from scRNA-seq data. Then, we wanted to further identify spatially variated immune cell subpopulations by combining both transcriptional and spatial information of each cell. Thus, a total of 2064 T cells were collected from melanoma scRNA-seq data^35^. Next, scSpace was utilized to recover the spatial characteristic within the scRNA-seq data referenced by spatial transcriptomics data derived from another experiment^36^.

As shown in Fig. 5a, T cells were classified into five refined subclusters by scSpace. Furthermore, the differential expression of these cell subsets was analyzed to retrieve marker genes for each subpopulation (Fig. 5b). Next, all T cells and malignant cells were distributed in the same pseudo-space (Fig. 5c), and the normalized pairwise distances between T cells in each subpopulation and malignant cells were calculated (Fig. 5d). The pairwise distances between malignant cells and T cells of C5, C1, C4, C2, and C3 subclusters in the pseudo-space increased gradually. Subsequently, we compared the two T cell subpopulations nearest (C5) and farthest (C3) from malignant cells in the melanoma microenvironment by differential expression analysis (Fig. 5e and Extended Data Fig. 5a). The result showed that IL7R, JUNB, TXNIP, DUSP1, and PLAC8 were highly expressed in C3, while TK1, AURKB, BIRK5, KIFC1, and MKI67 were specifically expressed in C5 (Fig. 5f and Extended Data Fig. 5b).

**Fig. 5.**
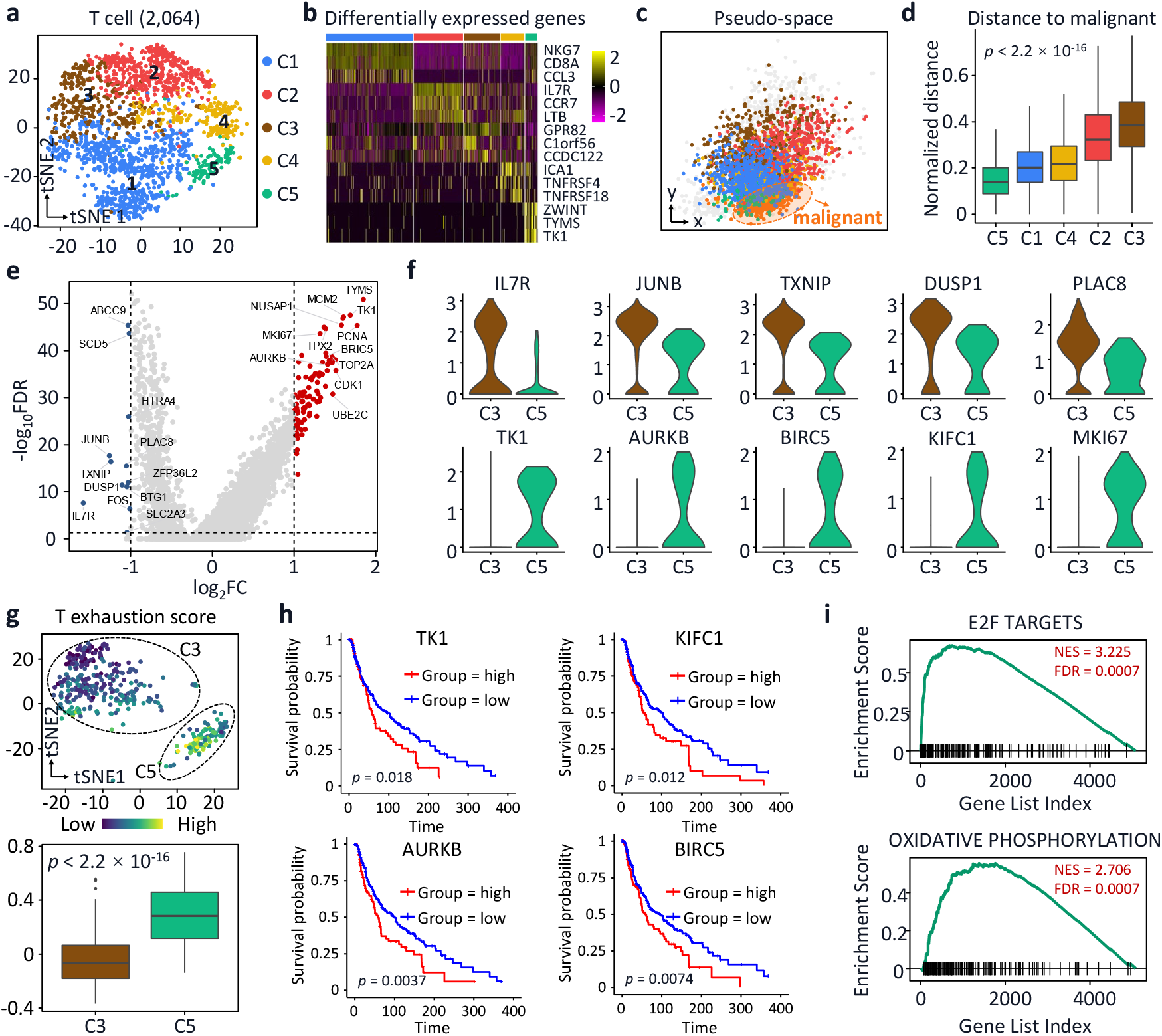
Spatial analysis of T cells subpopulations in melanoma. **a**, Space-informed classification of T cells generated five T cell subpopulations. **b**, Differential expression analysis of the five T cell subclusters. **c**, Pseudo-space of T cell subpopulations together with malignant cells constructed by scSpace. **d**, Normalized pairwise distances between malignant cells and cells in different subtypes. **e**, Differential expression analysis of C3 and C5, two subpopulations farthest and nearest from malignant cells. **f**, The expression of marker genes for C3 (Top) and C5 (Bottom) subpopulations. **g**, T exhaustion analysis of C3 and C5 subclusters. Top, tSNE layout of the two T cell subpopulations, C3 and C5. The color of each cell represents the TES according to the scalebar. Bottom, averaged TES for C3 and C5 subclusters (*p* < 2.2×^-16^). **h**, Survival probability analysis of the four highly expressed marker genes (TK1, KIFC1, AURKB, and BIRC5) of the C5 T cell subcluster. High expression of these genes exhibited a lower survival probability. **i**, Gene ontology analysis of marker genes enriched E2F targets and Oxidative phosphorylation pathways.

Further, we calculated the T exhaustion score (TES) for the two subpopulations using exhaustion-related genes (Extended Data Table S3). T cell exhaustion is the loss of T cell function in patients with common chronic infections and cancer. As a result of long-term exposure to persistent antigen and inflammation, exhausted T cells gradually lose their effective function. Our results showed that cells from C5 that were closer to malignant cells had higher TES (Fig. 5g), which is consistent with the T cell exhaustion hypothesis. We then performed a survival analysis of marker genes in the C5 subpopulation (Fig. 5h and Extended Data Fig. 5c). The results demonstrated that the high expression of these marker genes in T cells can significantly reduce the survival probability of patients, which is expected to become a therapeutic target for precision medicine in clinics. Indeed, therapeutic strategies targeting TK1^37^, KIFC1^38^, AURKB^39^, TPX2^40^, and BIRC5^41^ have been reported to be potentially effective. The gene set enrichment analysis enriched E2F targets, the oxidative phosphorylation, and another four pathways for C5 (Fig. 5i and Extended Data Fig. 5d), which are closely related to the tumor microenvironment.

### Reconstruction of cell pseudo-space captured invasion of myeloid subpopulations in Covid-19

We further examined whether scSpace could distinguish the spatial variation of cells between normal and disease conditions. Two scRNA-seq data^42^ with comparable expression states from a Covid-19 and a normal sample were collected to illustrate the performance of scSpace (Extended Data Fig. 6). By projecting single cells in normal and diseased tissues into the same pseudo-space with scSpace, we can compare the cell-type composition and proportion, the spatial distribution patterns, and the relative pairwise associations between cell subpopulations. More accurately describe the process of the occurrence and development of the disease, and find the key targets for the treatment of the disease.

After reconstructing spatial relationships of cells, the pseudo-space of the Control and Covid-19 group was established (Fig. 6a). We found a higher proportion of myeloid cells in the Covid-19 group than in the Control group, and they were closer to the epithelial cells (Fig. 6b). Consistently, myeloid cells were reported to infiltrate from the blood into the airway in several patients with Covid-19^43, 44^. As shown in Fig. 6c, myeloid cells were further divided into five subtypes, including diffusion component (DC), resident alveolar macrophages (AM), monocytes (Mon), monocyte-derived macrophages (MDM), and transitioning MDM (TMDM). The pairwise distances between these cell types and epithelial cells were significantly different in the Covid-19 group compared with the Control group (Fig. 6d). Notably, AM was predicted by scSpace to be the closest cell type to the epithelial cells, which is compatible with the common knowledge that AM is itself in the alveoli. Meanwhile, MDM/TMDM experienced the highest levels of invasion in Covid-19, consistent with previous research^44^. The two cell types that changed the most, MDM and TMDM, were selected for downstream analysis based on their dramatic decrease in the Covid-19 group versus the Control group (Fig. 6e).

**Fig. 6.**
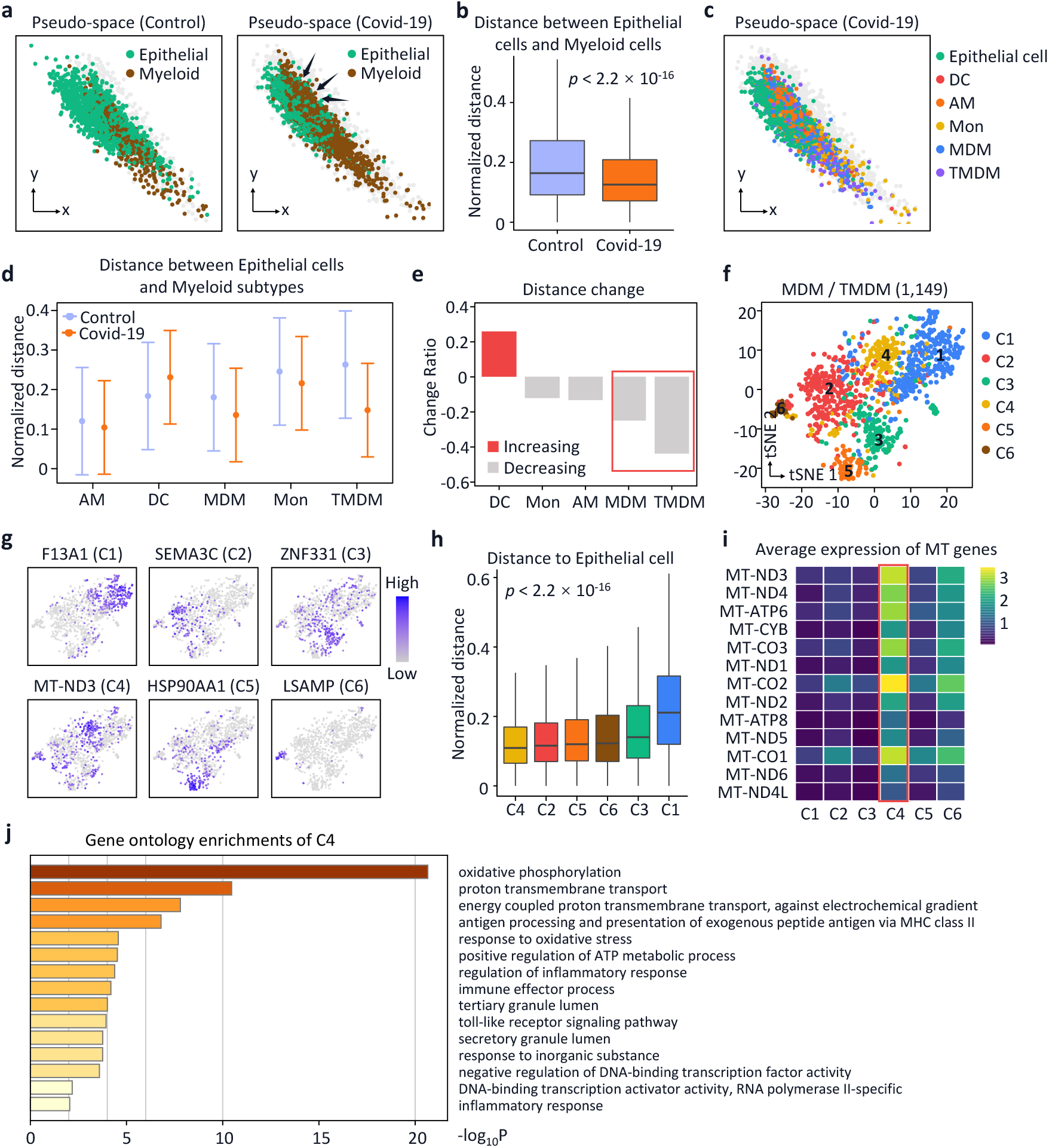
Spatial analysis of the invasion of myeloid subpopulations in Covid-19. **a**, The reconstructed pseudo-space for Control and Covid-19 group by scSpace. **b**, Normalized pairwise distances between epithelial cells and myeloid cells in Control and Covid-19 group. The distance in Covid-19 group was significantly lower (*p*<2.2×^-16^). **c**, The pseudo-space of epithelial cells and five subpopulations of myeloid cells. **d**, Normalized pairwise distances between epithelial cells and cells in different subtypes in Control and Covid-19 group. **e**, Distance changes in the Covid-19 group compared with the Control group. MDM and TMDM were dramatically decreased in the Covid-19 group. **f**, Refined classification of MDM and TMDM generated six subtypes. **g**, Expression value of marker genes in the six subpopulations of MDM and TMDM. **h**, Normalized pairwise distance between epithelial cells and six subpopulations of MDM and TMDM. **i**, Averaged expression level of mitochondrial-related genes in the six different subclusters of MDM and TMDM. **j**, Gene ontology enrichment of C4, the closest subpopulation to epithelial cells.

We thus clustered MDM and TMDM into six subpopulations by combining both spatial information and transcriptomes of each cell (Fig. 6f). The expression patterns of marker genes in each cell subpopulation indicated a strong spatial heterogeneity in MDM and TMDM (Fig. 6g). Then, we calculated the normalized pairwise distances between epithelial cells and different subpopulations and found that the C4 subcluster was nearest to epithelial cells (Fig. 6h). Compared with other cell subpopulations, C4 significantly overexpressed a large number of mitochondria-related genes (Fig. 6i) and subsequent gene ontology enrichment showed that these highly expressed genes were closely related to oxidative phosphorylation, proton transmembrane transport, energy metabolism, oxidative stress, etc. (Fig. 6j). These pathways played a crucial role in the infection process and had great therapeutic potential for the development of novel drugs to treat Covid-19^45^.

## Discussion

The spatial information is as important as transcriptional expression in the definition of cell identity. In brain tissue, for instance, cells with an identical cell type perform radically different functions in different regions^46^. The combination of the spatial and transcriptional characteristics of single cells will certainly make it possible to find more refined cell subpopulations. Although single-cell sequencing can investigate the molecular heterogeneity of cells at a high throughput and lead scientific research into the era of single-cell biology, due to technical limitations, it has to cause the loss of spatial information, which undoubtedly hinders the accurate description of cell identity. Meanwhile, spatial transcriptomics technologies such as ST, Visium, and Slide-seq, failed to analyze the differences between cells at single-cell resolution and other imaging-based methods, such as MERFISH, seqFISH, and STARmap, cannot achieve unbiased profiling of the whole transcriptome of individual cells.

On the other hand, *in silico* methods, such as NovoSpaRc^47^, Trendsceek^48^, and SpatialDE^49^ were designed to explore the spatial expression pattern of genes without further discussing the how the spatial heterogeneity could determine the cell type identification. SpaGCN^50^ and STAGATE^51^ enabled the identification of spatial domains with coherent expression and histology but didn’t attach the importance of cell independence. Therefore, inferring the spatial information within scRNA-seq data is crucial to further reveal the identity of individual cells.

Here, we used a transfer learning model to integrate single-cell and spatial transcriptomics to retrieve the spatial relationship of cells. Our results demonstrated that spatial information could indeed improve the accuracy and fineness of cell clustering no matter in simulated or biological datasets. Finally, we applied scSpace to multiple circumstances, such as melanoma and Covid-19. As a result of long-term exposure to persistent antigen and inflammation, exhausted T cells gradually lose effector function and memory T cell features begin to be lost. Spatial analysis of T cell subpopulations in melanoma revealed T cell exhaustion happened near the cancer microenvironment, which indicated that scSpace has potential to determine tumor stages by distinguishing the nearest subgroup of T cells and calculating its TES. The pandemic of Covid-19 make it particularly important for comprehensive understanding of its pathogenesis. The use of scSpace has provided spatial therapeutic markers and brought new insight into the study of Covid-19.

## Acknowledgement

This work is supported by National Natural Science Foundation of China (81973701), the Natural Science Foundation of Zhejiang Province (LZ20H290002), and Alibaba Cloud provided by Alibaba-Zhejiang University Joint Research Center of Future Digital Healthcare.

## Author contributions

X. F., B. Z., and J. L. conceived the study. J. L., J. Q., Z. L., and Y. C. collected datasets involved in this article, benchmarked all methods, and participated in the development of the scSpace algorithm. Y. Z. analyzed and the biological meaning of the results predicted by scSpace. X. S., J. C., Y. C., W. G., P. Y., Y. H., H. B., Q. C., and M. L. provided a lot of advice on algorithm implementation and biological applications. W. G., Y. H., and H. B. provided valuable advice on gene ontology analysis in this study. All authors wrote the manuscript, read and approved the final manuscript.

## Competing interests

The authors declare no competing interests.

## Methods

### Data processing

We applied the Seurat^52^ R package (v4.1.0) to pre-process the scRNA-seq and spatial transcriptomics data used in scSpace. We first filtered low-quality cells and genes following the standard quality control step of scRNA-seq data analysis. Next, the raw count data were normalized using the ‘NormalizeData’ function with the default parameters, and 2000 (by default) highly variable genes were selected using the ‘FindVariableFeatures’ function with the ‘vst’ method.

### Identification of spatially associated subpopulations by scSpace

The workflow of scSpace is shown in Fig. 1a and Extended Data Fig. 1a, including latent biological feature representation extraction, spatial reconstruction, and space-informed cell clustering in three steps.

#### Latent biological feature representation extraction

Using the scRNA-seq data *X*_*S*_ and a spatial transcriptomics reference *X*_*T*_ as inputs, scSpace first applied a transfer learning method^53^ TCA to eliminate the batch effect of these two types of data and extract a good feature representation across these two domains with true biological characteristics.

Denote *P*(*X*_*S*_) and *Q*(*X*_*T*_) as the marginal distributions of scRNA-seq data and spatial transcriptomics data, respectively. We assume that *P*(*X*_*S*_) ≠ *Q*(*X*_*T*_), but there exists a transformation *ϕ* such that *P*(*ϕ*(*X*_*S*_)) ≈ *P*(*ϕ*(*X*_*T*_)). Especially, the transformation *ϕ* satisfies, 1) the distance between *P*(*ϕ*(*X*_*S*_)) and *P*(*ϕ*(*X*_*T*_)) is small, and 2) *ϕ*(*X*_*S*_) and *ϕ*(*X*_*T*_) preserve important properties of *X*_*S*_ and *X*_*T*_.

Next, the Maximum Mean Discrepancy (MMD) distance between these two distributions is calculated following,

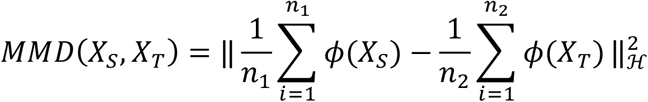

Where ‖ · ‖_ℋ_ is the Reproducing Kernel Hilbert Space (RKHS) norm^54^, and the transformation *ϕ* can be found by minimizing the distance.

In order to effectively find *ϕ*, a kernel matrix *K* is introduced:

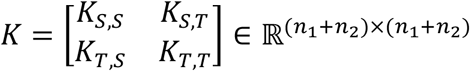

And the MMD distance can be presented as:

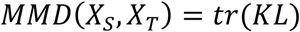

Where:

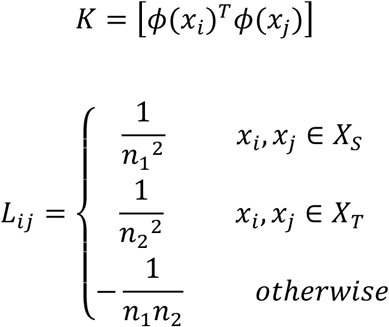

Next, the empirical kernel map step is applied by decomposing the kernel matrix

*K* as *K* = (*KK*^−1/2^)(*K*^−1/2^*K*), and then transforms this to an *m*-dimensional space using the matrix 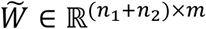, and the kernel matrix is:

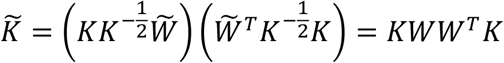

Where 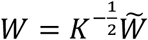. Then, the MMD distance can be defined as:

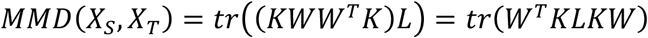

Finally, objective function is defined as:

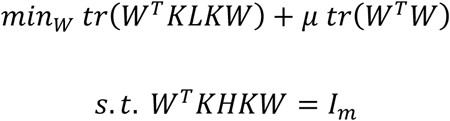

Where *μ* > 0 is a tradeoff parameter, 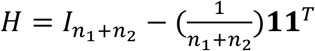 is the centering matrix, and *I*_*m*_ ∈ ℝ^*m*×*m*^ is the identity matrix. The objective function can be reformulated as:

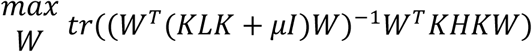

And the W solution are the *m* leading eigenvectors of (*I* + *μKLK*)^−1^*KHK*, i.e., the latent feature representation across scRNA-seq data and spatial transcriptomics data with true biological characteristics.

#### Spatial reconstruction

Once the latent biological feature representation across scRNA-seq and spatial transcriptomic data are extracted, a multi-layer fully connected neural network model is applied to spatial reconstruction of scRNA-seq data. The size of input layer of this model is equal to the dimension of the latent biological feature representation, and the size of output layer is corresponding to the dimension of spatial coordinates. Denote 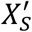 and 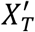are latent biological feature representation of scRNA-seq data and spatial transcriptomics data, respectively. We assume that the spatial locations of cells are related to their latent biological feature representation:

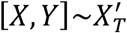

Where *X* and *Y* are spatial coordinates of cells (or spots) in spatial transcriptomics data 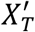. scSpace first trains the model on scRNA-seq data using mean squared error (MSE) loss function. Once training is finished, scSpace then applies the model to scRNA-seq data 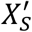, and the spatial information of each single cell is reconstructed (we term ‘pseudo-space’).

#### Space-informed clustering

scSpace applies space-informed clustering to identify spatially heterogeneous single-cell subclusters based on gene expression and pseudo-space information of cells in scRNA-seq data. In detail, a gene expression graph *G*_*g*_(*V, E*_1_) and a space graph *G*_*S*_(*V, E*_2_) are constructed respectively using *k*-nearest neighbor (KNN) algorithm. Since our goal is to find spatially heterogeneous subclusters that may be similar in gene expression, the space graph *G*_*S*_(*V, E*_2_) is then transformed to the spatial weight *w* of each edge in gene expression graph *G*_*g*_(*V, E*_1_):

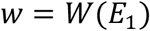

Finally, scSpace applies unsupervised clustering on spatial-weighted gene expression graph using Leiden algorithm^55^.

### Simulated data analysis

We applied Splatter R package (v1.16.1) ^56^ to simulate 50 paired scRNA-seq and spatial transcriptomics data with 5000 expression genes (Extended Data Fig. 1b). In order to simulate the original batch effect between these two types of data, we set up 2 batches representing scRNA-seq and spatial transcriptomics data, respectively. Moreover, for the robustness of results, we set a gradient from 500 to 2000 for the number of cells, and a gradient from 3 to 8 for the number of cell populations.

Next, for each cell, a pseudo spatial coordinate was assigned based on random sampling and normal distribution strategy. We randomly choose 2 to 4 cell populations and set the probabilities of a gene being differentially expressed in each of them as 0.01 (0.1 ∼ 0.2 for remaining cell populations by contrast). Then, we assigned these cell populations as different spatially associated subpopulations with similar transcriptome.

We benchmarked scSpace with 3 classical clustering algorithms: Louvain, K-means and Hierarchical clustering. We used adjusted rand index (ARI) and normalized mutual information (NMI) measurements to evaluate the performance of clustering result. We also performed Pearson correlation analysis to evaluate the spatial reconstruction results of scSpace.

### Validation of scSpace on spatial transcriptomics data

*Human dorsolateral prefrontal cortex (DLPFC) data*. The human DLPFC spatial transcriptomics data ^31^ (Visium, 10x Genomics) was downloaded from http://research.libd.org/spatialLIBd. We selected slice 151674 as spatial transcriptomics data reference, and other 11 slices as simulated scRNA-seq data. The distance between cells were normalized with the following formula:

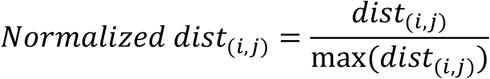

The differential expression genes between cortex layers were calculated by the function ‘FindAllMarkers’ of Seurat R package (v4.1.0) with the default two-tailed Wilcoxon rank sum test.

*Mouse V1 neocortex data*. The mouse V1 neocortex spatial transcriptomics data (STARmap)^9^ was downloaded from https://www.starmapresources.org/data. We

selected two replicates with 1020 genes as spatial transcriptomics data reference and simulated scRNA-seq data, respectively.

### Validation of scSpace on scRNA-seq data

#### Mouse V1 neocortex data

The mouse V1 neocortex scRNA-seq data were downloaded from the RNA-seq data repository of Allen Brain Atlas website (https://portal.brain-map.org/atlases-and-data/rnaseq/mouse-v1-and-alm-smart-seq) and then we down sampled to 5000 cells from the data. The mouse V1 neocortex STARmap data were utilized as spatial references. The differential expression genes between cortex layers were calculated using the ‘FindAllMarkers’ function in Seurat R package (v4.1.0).

#### Mouse intestines data

The mouse intestine scRNA-seq data^32^ were downloaded from the Gene Expression Omnibus (GEO, GSE109413). We also downloaded another set of spatial transcriptomics data^57^ (GSE169749) as the spatial reference. The distance between cells were normalized as described before.

#### Mouse liver data

The mouse liver scRNA-seq data^33^ and the spatial transcriptomics reference^58^ were obtained from GEO (GSE84498) and the Liver Cell Atlas (http://www.livercellatlas.org), respectively. The distance between cells were normalized as described before.

#### Human developing heart dat

The human developing heart scRNA-seq data and the spatial transcriptomics reference data sequenced by Asp ^34^ et al. were downloaded from https://www.spatialresearch.org. We used Louvain as benchmarking clustering method compared for scSpace. Statistical differences in the distance between cells of different subpopulations were assessed with Mann-Whitney test. The differential expression genes between different cell subpopulations were calculated by Seurat R package (v4.1.0).

### Discovery of fine subpopulations in human cortex scRNA-seq data

For the human cortex scRNA-seq data (SMART-Seq v4), we downloaded it from Allen Brain Atlas (https://portal.brain-map.org/atlases-and-data/rnaseq/human-multiple-cortical-areas-smart-seq) and then down sampled to 4000 cells. The human DLPFC data (slice 151674) was applied as spatial reference. We extracted C1, C2, C3, C4, and C9 clusters for IT subpopulation analysis. Layer proportion and normalized distance to L1/WM of every cluster were calculated. We further investigated the specifically expression genes of each IT subpopulation using Seurat R package (v4.1.0). The *in situ* hybridization (ISH) images from visual cortex (LAMP5, GRIK1) or temporal cortex (GABRG1, PCP4, and RXFP1) of adult human brain were downloaded from Allen Human Brain Atlas: http://human.brain-map.org/.

### Spatial analysis of T cells subpopulations in melanoma

The human melanoma scRNA-seq^35^ and spatial transcriptomics data^36^ were accessed from GEO (GSE72056) and https://www.spatialresearch.org, respectively. After assigning every single cell a pseudo-space coordinates, 2064 T cells were extracted for spatial-informed sub-clustering, and We calculated the normalized distance of every T subpopulation to malignant cell. We then selected subpopulation C5 closest to malignant cell and subpopulation C3 farthest to malignant cell for differential expression gene analysis using the ‘FindMarker’ function of Seurat R package (v4.1.0). To characterize T cell exhaustion of C3 and C5, we used a set of exhaustion gene signature from Zheng^59^ et.al. And the average expression levels of exhaustion gene signatures were calculated using the ‘AddModuleScore’ function of Seurat R package (v4.1.0).

To further examine the clinical relevance and biological function of C5, we performed some survival analysis and pathway enrichment analysis. For survival analysis, RNA-seq and clinical data of melanoma patients (cancer study id: skcm tcga) were obtained from TCGA using cgdsr R package (v1.3.0). The samples were divided into two groups along with low (25%) and high (75%) target genes expression for all patients, and then survival curves of these two groups of patients were estimated by the Kaplan–Meier method using survival R package (v3.2-13). For pathway enrichment analysis, the hallmark gene sets were downloaded from the Molecular Signatures Database^59^ (MSigDB v7.4) using msigdbr R package (v7.4.1) and gene set enrichment analysis was performed^60^ fgsea R package (v1.18.0).

### Spatial analysis of the invasion of myeloid subpopulations in Covid-19

The molecular single-cell lung atlas of lethal Covid-19 by Melms^42^ et.al. was downloaded from https://singlecell.broadinstitute.org/single_cell/study/SCP1219, and we randomly down sampled to 10000 cells. As a spatial reference, the normal human lung spatial transcriptomics data by Lakshminarasimha et.al. was downloaded from GEO (GSE178361). The change ratio of normalized distance between epithelial cell and each myeloid subtype in Covid-19 group and control group were calculated later.

A total of 1149 monocyte-derived macrophages (MDMs) and transitioning MDMs (TMDMs) were extracted for next space-informed clustering. We then calculated the normalized distance between every MDM/TMDM subpopulation and epithelial cell to evaluate the degree of infiltration. For the subpopulation C4 whose cells were closest to the epithelial cells in the pseudo-space, we performed the pathway enrichment analysis using the Metascape^61^ (https://metascape.org) to investigate the biological functions.

## Data availability

Data used in this study were downloaded from publicly available datasets. No experimental data conducted by ourselves were used.

## Code availability

The scSpace algorithm and related analysis is available at GitHub: https://github.com/ZJUFanLab/scSpace.

## Statistics

Python (version 3.8.5) and R (version 4.1.0) were used for the statistical analysis.

## Notes

### Competing Interest Statement

The authors have declared no competing interest.

