## Supplemental information for "Reconstruction of the cell pseudo-space from single-cell RNA sequencing data with scSpace"

#### Supplementary Tables

**Table S1. Benchmark for the evaluation of scSpace versus other methods using simulated data.** ARI, Adjusted rand index. NMI, normalized mutual information.

| Simulations | Subcluster numbers | Methods | ARIs (all) | NMIs (all) | ARIs (subcluster) | NMIs (subcluster) |
|---|---|---|---|---|---|---|
| Simulation 1 to Simulation 15 | 2 | **scSpace** | **0.983±0.032** | **0.960±0.027** | **0.875±0.113** | **0.783±0.126** |
| | | Louvain | 0.778±0.184 | 0.817±0.110 | 0.000±0.214 | 0.001±0.174 |
| | | Kmeans | 0.766±0.187 | 0.854±0.101 | 0.000±0.311 | 0.000±0.260 |
| | | Hclust | 0.502±0.368 | 0.610±0.364 | 0.000±0.012 | 0.005±0.007 |
| Simulation 16 to Simulation 35 | 3 | **scSpace** | **0.905±0.111** | **0.888±0.063** | **0.720±0.106** | **0.639±0.108** |
| | | Louvain | 0.705±0.145 | 0.794±0.082 | 0.379±0.207 | 0.369±0.189 |
| | | Kmeans | 0.665±0.143 | 0.805±0.083 | 0.400±0.233 | 0.437±0.215 |
| | | Hclust | 0.545±0.175 | 0.671±0.167 | 0.002±0.010 | 0.011±0.008 |
| Simulation 36 to Simulation 50 | 4 | **scSpace** | **0.858±0.121** | **0.840±0.077** | **0.649±0.148** | **0.657±0.145** |
| | | Louvain | 0.691±0.172 | 0.729±0.108 | 0.381±0.209 | 0.383±0.192 |
| | | Kmeans | 0.691±0.168 | 0.728±0.097 | 0.398±0.220 | 0.390±0.217 |
| | | Hclust | 0.342±0.213 | 0.543±0.258 | 0.001±0.006 | 0.013±0.011 |

**Table S2. *Pearson* correlation of the Pseudo-space generated by scSpace and ground truth using benchmark datasets.**

| Simulation | Subcluster number | Method | Pseudo-space *Pearson* correlation |
|---|---|---|---|
| Simulation 1 to 15 | 2 | scSpace | 0.959 ± 0.041 |
| Simulation 16 to 35 | 3 | | 0.937 ± 0.104 |
| Simulation 36 to 50 | 4 | | 0.919 ± 0.068 |

**Table S3. Exhaustion-related genes of T cells.** Total 74 genes listed.

| Gene ID | Gene Symbol | Gene ID | Gene Symbol |
|---|---|---|---|
| 84868 | HAVCR2 | 3002 | GZMB |
| 5133 | PDCD1 | 1390 | CREM |
| 953 | ENTPD1 | 3108 | HLA-DMA |
| 3604 | TNFRSF9 | 639 | PRDM1 |
| 6348 | CCL3 | 5315 | PKM |
| 22822 | PHLDA1 | 4643 | MYO1E |
| 55423 | SIRPG | 2530 | FUT8 |
| 1493 | CTLA4 | 1846 | DUSP4 |
| 201633 | TIGIT | 5873 | RAB27A |
| 116841 | SNAP47 | 3122 | HLA-DRA |
| 939 | CD27 | 79713 | IGFLR1 |
| 5996 | RGS1 | 10563 | CXCL13 |
| 7133 | TNFRSF1B | 3458 | IFNG |
| 54 | ACP5 | 140739 | UBE2F |
| 84632 | AFAP1L2 | 9452 | ITM2A |
| 4647 | MYO7A | 3399 | ID3 |
| 9495 | AKAP5 | 10421 | CD2BP2 |
| 9289 | ADGRG1 | 55501 | CHST12 |
| 9760 | TOX | 1509 | CTSD |
| 143903 | LAYN | 6774 | STAT3 |
| 952 | CD38 | 684 | BST2 |
| 3682 | ITGAE | 10663 | CXCR6 |
| 5997 | RGS2 | 5900 | RALGDS |
| 894 | CCND2 | 7412 | VCAM1 |
| 2280 | FKBP1A | 10906 | TRAFD1 |
| 4522 | MTHFD1 | 9144 | SYNGR2 |
| 130589 | GALM | 9218 | VAPA |
| 1435 | CSF1 | 3430 | IFI35 |
| 51429 | SNX9 | 967 | CD63 |
| 29851 | ICOS | 4664 | NAB1 |
| 1130 | LYST | 11315 | PARK7 |
| 114614 | MIR155HG | 5573 | PRKAR1A |
| 7167 | TPI1 | 3560 | IL2RB |
| 1757 | SARDH | 9324 | HMGN3 |
| 25824 | PRDX5 | 10797 | MTHFD2 |
| 3902 | LAG3 | 54602 | NDFIP2 |
| 64231 | MS4A6A | 5551 | PRF1 |

### Supplementary Figures

#### Extended Data Fig 1. Detailed workflow of scSpace and performance evaluations on simulated datasets.

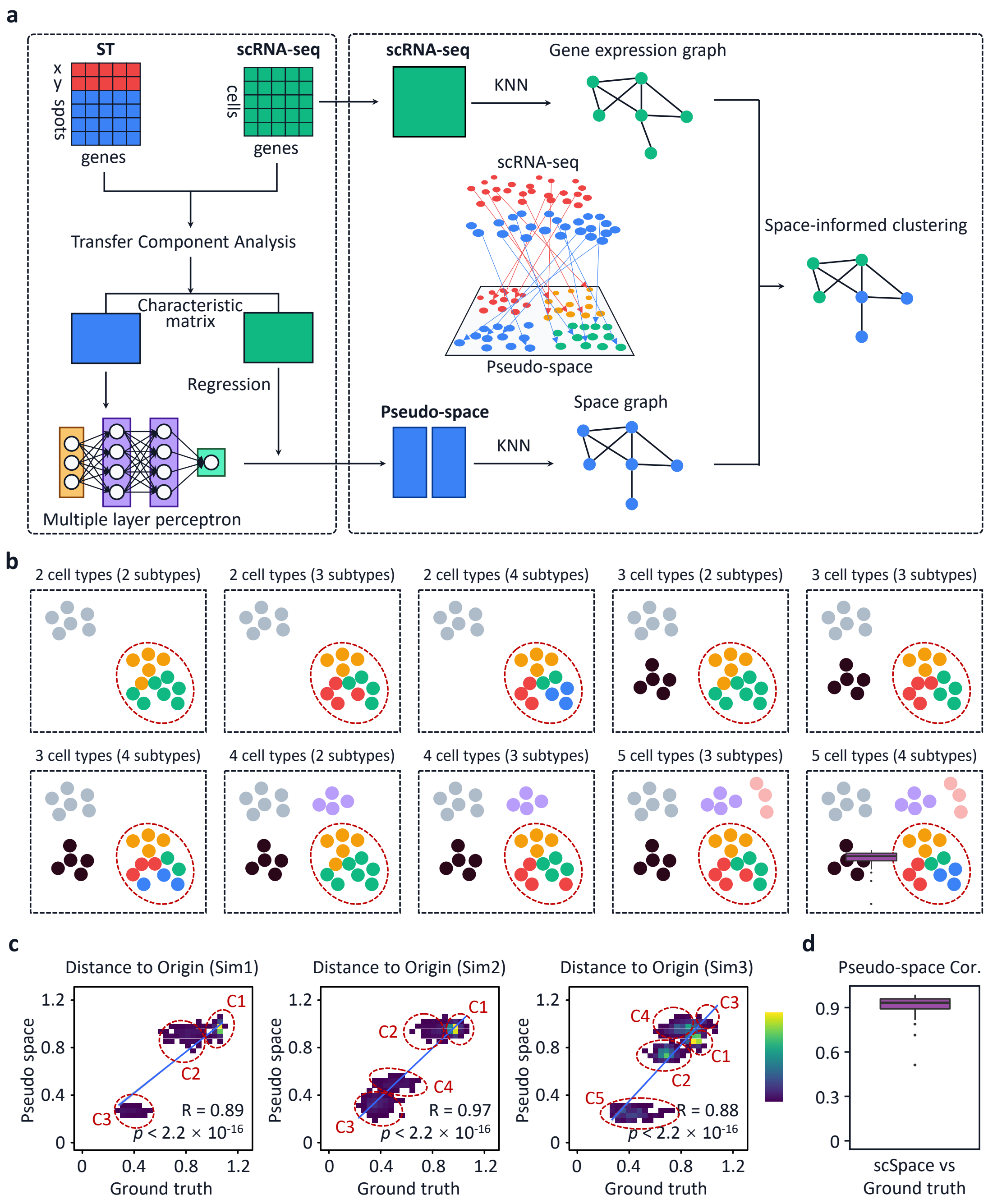


**a**, Integration procedure of scSpace. Spatially resolved transcriptomics and scRNA-seq data matrixes were taken as input. Then, Transfer Component Analysis (TCA) was conducted to extract each characteristic matrix from both data. A multiple layer perceptron (MLP) model was used to combine the two matrixes for the pseudo-space reconstruction. KNN algorithm was used to generate the gene expression graph and the space graph from scRNA-seq and pseudo-space data, respectively. Finally, spatially variated subclusters were obtained by the space-informed clustering.

**b**, Simulation of spatially resolved transcriptomics data. 50 paired scRNA-seq and spatial transcriptomics data with 5000 expression genes were simulated to evaluate the performance of scSpace. We examined whether scSpace could distinguish cell subtypes with or without spatial heterogeneity. **c**, Pairwise distance between cells in the pseudo-space and the ground truth for the three examples shown in Fig.1b of the manuscript. **d**, Pearson correlation of all simulations between scSpace and the ground truth.

#### Extended Data Fig 2. Reconstructing the hierarchical structure of human brain cortex using spatial transcriptomics data.

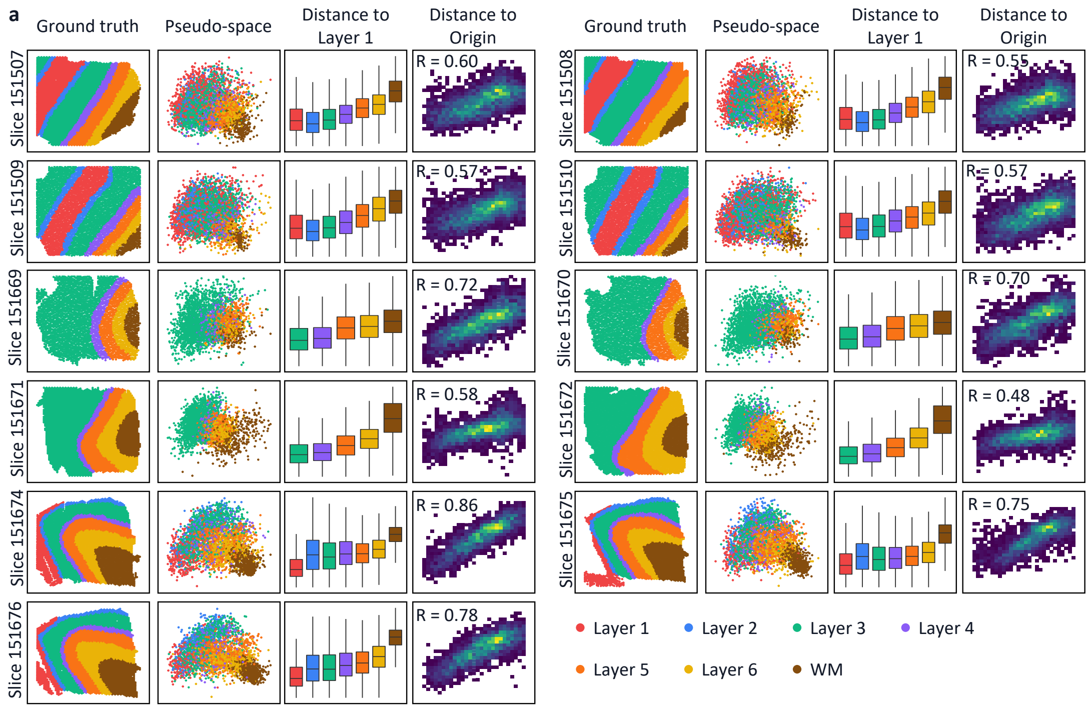


**a**, Spatial reconstruction of human DLPFC layers using one slice as the spatial reference and another as the test set by removing the cell coordinate information. Extra 11 examples in addition to Figure 2 in the manuscript were showed. Each slice was consisted of four figures. First, the annotation of each spot in the ground truth. Second, the pseudo-space generated by scSpace. Third, averaged pairwise distances between cells from different layers and Layer 1. Fourth, Pairwise distance between spots in the pseudo-space and the ground truth.

**Extended Data Fig 3. Reconstructing the zonation of human liver lobules using regional scRNA-seq data.**

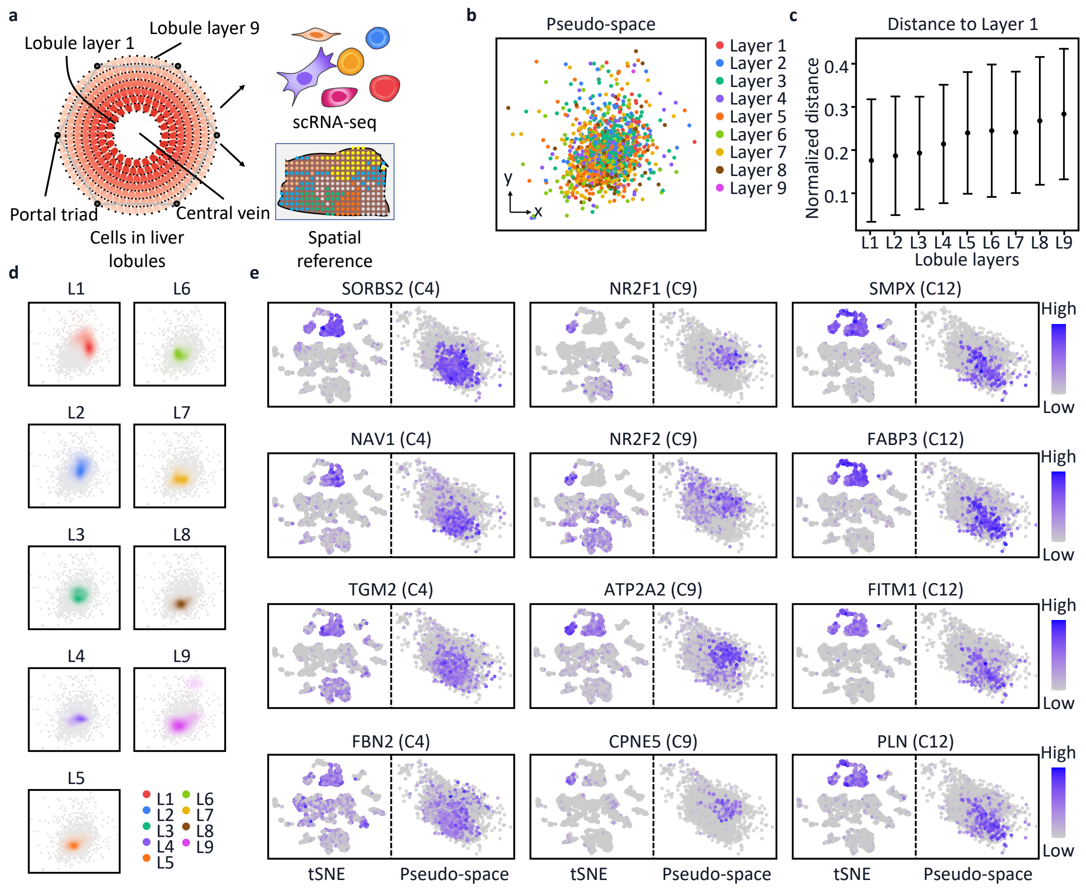


**a**, Schematic of the spatial reconstruction of liver lobules using biological scRNA-seq data from different layers. Both the single-cell data and spatial reference were accessed from experimental data. **b**, The pseudo-space of liver cells from nine layers generated by scSpace. **c**, Average pairwise distances between cells in different layers and cells in the Layer 1 from the near to the distant. **d**, Spatial distribution density of cells in each liver lobule layer. **e**, Supplementary information for Figure 3i in the manuscript. Expression patterns of marker genes for C4, C9, and C12 subpopulations in the pseudo-space, respectively.

**Extended Data Fig 4. Detailed differential expression analysis results.**

**a**

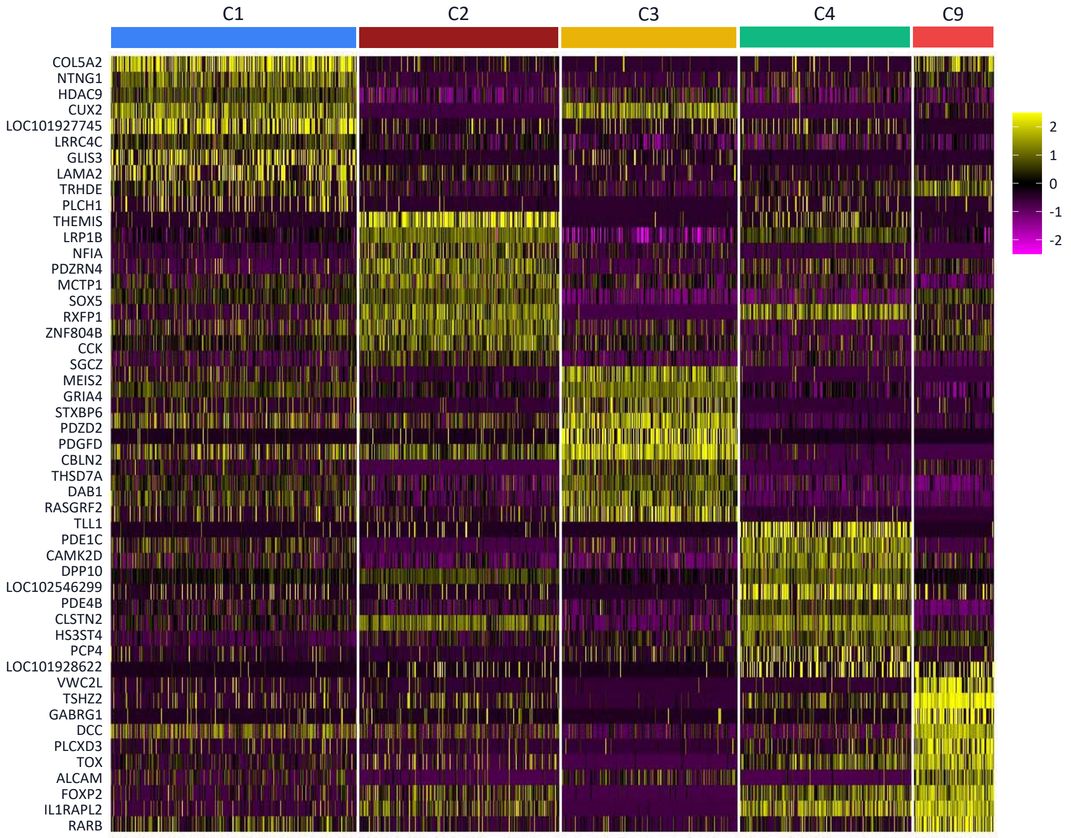


**a**, Differential expression analysis of the five subpopulations (C1, C2, C3, C4, and C9) of intratelencephalic (IT) neurons using both the transcriptional and spatial information by scSpace.

**Extended Data Fig 5. Supplementary information for the space-informed clustering of T cells in melanoma.**

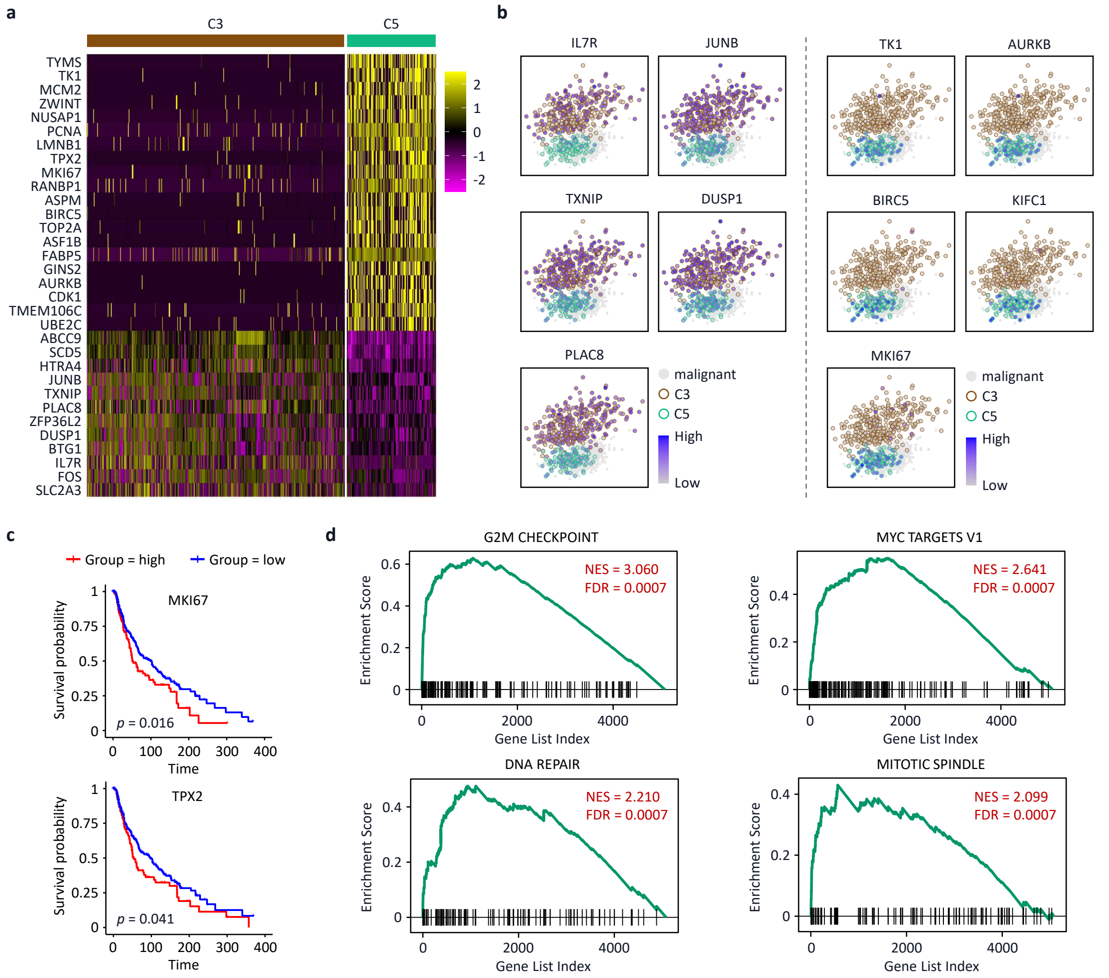


**a**, Differential expression analysis of C3 and C5 subclusters. **b**, Expression of C5 and C3 marker genes in the pseudo-space (IL7R, JUNB, TXNIP, DUSP1, and PLAC8 for C3, and TK1, AURKB, BIRC5, KIFC1, and MKI67 for C5 ) and their distance from malignant cells. **c**, Survival probability analysis of another two highly expressed marker genes (MKI67 and TPX2) of the C5 T cell subcluster. **d**, Gene set enrichment analysis of marker genes enriched G2M checkpoint, MYC targets V1, DNA repair, and mitotic spindle pathways.

**Extended Data Fig 6. Comparison of single-cell RNA-seq data between Covid-19 and Control Groups.**

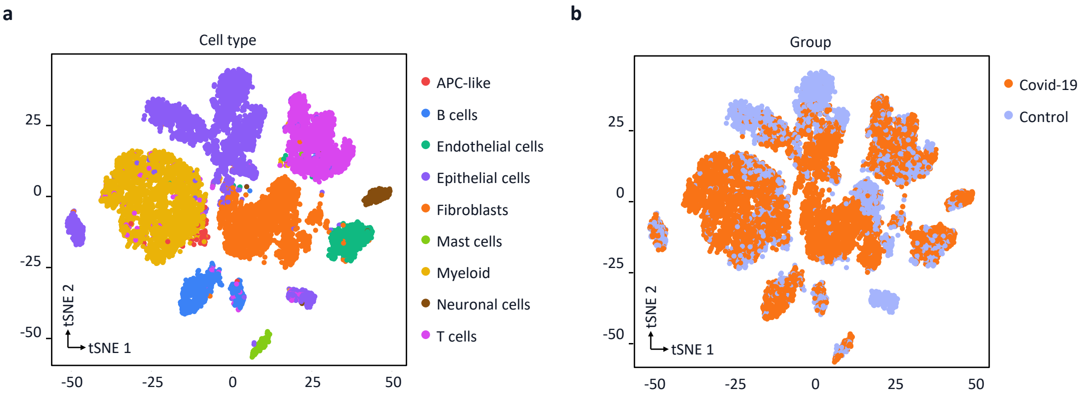


**a**, Joint analysis for clustering and annotation of single-cell RNA-seq data in Covid-19 and Control groups. **b**, Covid-19 group and Control group share comparable cell-state space in the gene expression of individual cells.
